# Microbubble Track-based Functional Ultrasound Localization Microscopy in Awake Mice

**DOI:** 10.1101/2025.08.18.670859

**Authors:** Yike Wang, Matthew R. Lowerison, Zhe Huang, YiRang Shin, Bing-Ze Lin, Pengfei Song

**Affiliations:** Department of Biomedical Engineering, Duke University, Durham, NC 27708 USA; Department of Electrical and Computer Engineering, University of Illinois Urbana–Champaign, Urbana, IL 61820 USA

**Keywords:** Functional ultrasound imaging, ultrasound localization microscopy, awake animal neuroimaging, cerebral blood flow

## Abstract

Functional neuroimaging with ultrafast ultrasound is an emerging neuroimaging tool for studying neural activities in the rodent brain. Existing methods, however, are challenged by the compromise between functional imaging sensitivity (i.e., sensitivity in detecting neural responses) and spatial resolution. For example, functional ultrasound (fUS) uses native red blood cells (RBCs) as imaging targets, which offers high functional imaging sensitivity but limited spatial resolution that is confined by the diffraction limit of ultrasound. On the other hand, functional ultrasound localization microscopy (fULM) employs intravenously injected microbubble (MB) as contrast agent to achieve super-resolved spatial resolution but at the cost of functional imaging sensitivity. This study aims to address this challenge by developing a novel, MB track-based hemodynamic activity estimation method to enhance the functional imaging sensitivity of fULM. Our approach involves conducting functional correlation analysis using the MB signals acquired from the entire MB movement track rather than individual MB centroid locations, which overcomes the signal sparsity issue in fULM. To further boost the functional sensitivity of fULM, we developed a novel approach based on indwelling jugular vein catheters to achieve fULM imaging in awake mice. The *in vivo* imaging results demonstrate that the proposed techniques successfully enhanced the functional imaging sensitivity of fULM without compromising its high spatial resolution. In the whisker stimulation experiments, the proposed technique enabled detection of significantly activated brain regions within fewer than five stimulation cycles (5 minutes of acquisition), reducing the required time by over 50% compared to conventional fULM.

## I. INTRODUCTION

**F**unctional ultrasound (fUS) has been rapidly evolving over the past decade as a powerful neuroimaging technique in neuroscience [1], [2]. Leveraging the high temporal resolution of ultrafast ultrasound [3], fUS captures cerebral blood flow (CBF) dynamics and infers neuronal activity through neurovascular coupling [4]. Compared to traditional neuroimaging techniques, fUS offers a balanced spatiotemporal resolution and imaging depth. For example, while functional magnetic resonance imaging (fMRI) enables whole-brain coverage, it suffers from relatively low spatial resolution for resolving fine-scale vascular or neural structures. Optical imaging methods, although capable of achieving high spatial resolution, are limited to superficial cortical layers with a noninvasive imaging setup, or limited field-of-view (FOV) with fiber optic implants. In contrast, fUS has better imaging coverage and depth of penetration than optical methods, and better spatial resolution (∼100 µm for 15 MHz) than fMRI [5], [6], [7]. In addition to the balanced imaging characteristics, fUS can also be performed in awake and freely moving animals [8], [9], [10], [11]. This is a notable advantage over techniques like fMRI that often require anesthesia or physical restraint.

Despite these strengths, fUS is still largely a mesoscopic imaging technique that does not provide microscopic insights offered by optical imaging techniques. For example, neural activity can induce subtle and spatially heterogeneous and local changes in CBF that reflect the function of small-scale microcircuits [12], [13], [14]. These patterns are challenging to resolve with conventional fUS [15]. Additionally, neurovascular coupling primarily occurs in the intraparenchymal vessels with diameters smaller than 30 μm [4], [16], which is beyond the resolution limit of fUS (i.e., ∼100 μm at 15 MHz). Recently, the emergence of functional ultrasound localization microscopy (fULM) addressed this limitation of fUS [16]. fULM was built on the foundation of ultrasound localization microscopy (ULM) [17], [18], [19], [20], [21], where intravenously injected microbubbles (MBs) are localized and tracked to achieve super-resolution imaging. fULM adapts this approach to capture microvascular hemodynamics by analyzing the fluctuations in microbubble (MB) counts over time, which reflects CBF variations that are correlated with neural activities. Importantly, fULM improves the spatial resolution by approximately 10-fold over fUS while preserving the same imaging depth of penetration and FOV [16].

The compromise that fULM makes to gain in spatial resolution is the reduction of sensitivity in detecting neural responses that are reflected in CBF variations. Unlike Fus which *directly* measures CBF using the backscattered red blood cell (RBC) signals, fULM *indirectly* measures CBF by estimating the MB count and its fluctuations in time. This makes the fULM measurements highly dependent on the local MB concentration in the blood stream and the localization and tracking algorithms used for MB count estimation. As a result, fULM requires substantially longer acquisition time with repeated stimulation cycles to average out the uncertainties associated with MB count (i.e., baseline MB fluctuations) and boost the functional imaging sensitivity to CBF variations. For instance, in the whisker stimulation regime, fUS only needs a few stimulation cycles (typically less than 5) in anesthetized animals and a single cycle in awake mice [7], but fULM typically requires over 10 cycles of stimulations to accumulate adequate signal and average out the baseline noise to detect cortical and thalamic responses [16]. Such long data acquisition time and the need of repeated stimulation greatly reduce the temporal resolution and the practicality of fULM as a useful and viable neuroimaging tool.

Recent advancements aim to mitigate this limitation. For example, deep-learning-based algorithms have been developed to enhance MB localization and tracking, enabling fULM imaging at higher MB concentrations [22]. MB backscattering intensity signal was also proposed to be combined with MB count signal to boost fULM sensitivity *in vivo* [23]. Although these studies have improved the efficiency of fULM, its functional sensitivity still remains substantially lower than that of conventional fUS imaging. Therefore, further enhancing fULM sensitivity remains a critical challenge. To this end, here we propose a novel MB track-based correlation analysis method to improve the functional sensitivity of fULM with no compromise to its spatial resolution. In addition, we developed a novel awake imaging setup for robust fULM. A jugular vein catheterization approach has been adopted to ensure stable MB delivery during imaging, which allows robust detection of whisker-evoked brain activation using fewer stimulation cycles than conventional fULM.

The remainder of this paper is organized as follows: Section II introduces the principles of the proposed track-based analysis method and compares it with conventional pixel-based fULM approaches. Section III details the material and methods. Section IV presents the results from both *in silico* validation with anesthetized rats and *in vivo* imaging experiments with awake mice. Finally, Section V concludes the study and discusses the limitations and directions for future research.

## II. Principles of MB-Track-based fULM

To facilitate the description of the principles of MB track-based fULM, we constructed a controlled simulation based on a single, targeted vessel (Fig. 1a), which was randomly selected from an *in vivo* fULM dataset that was previously published in [22]. The purpose of using *in vivo* dataset was to obtain anatomically realistic vascular structures. However, no functional-related information from the dataset was utilized. Instead, we manually defined flow changes to simulate a well-controlled functional activation process, which was modulated as variations in the number of detected MB tracks.

**Fig. 1.**
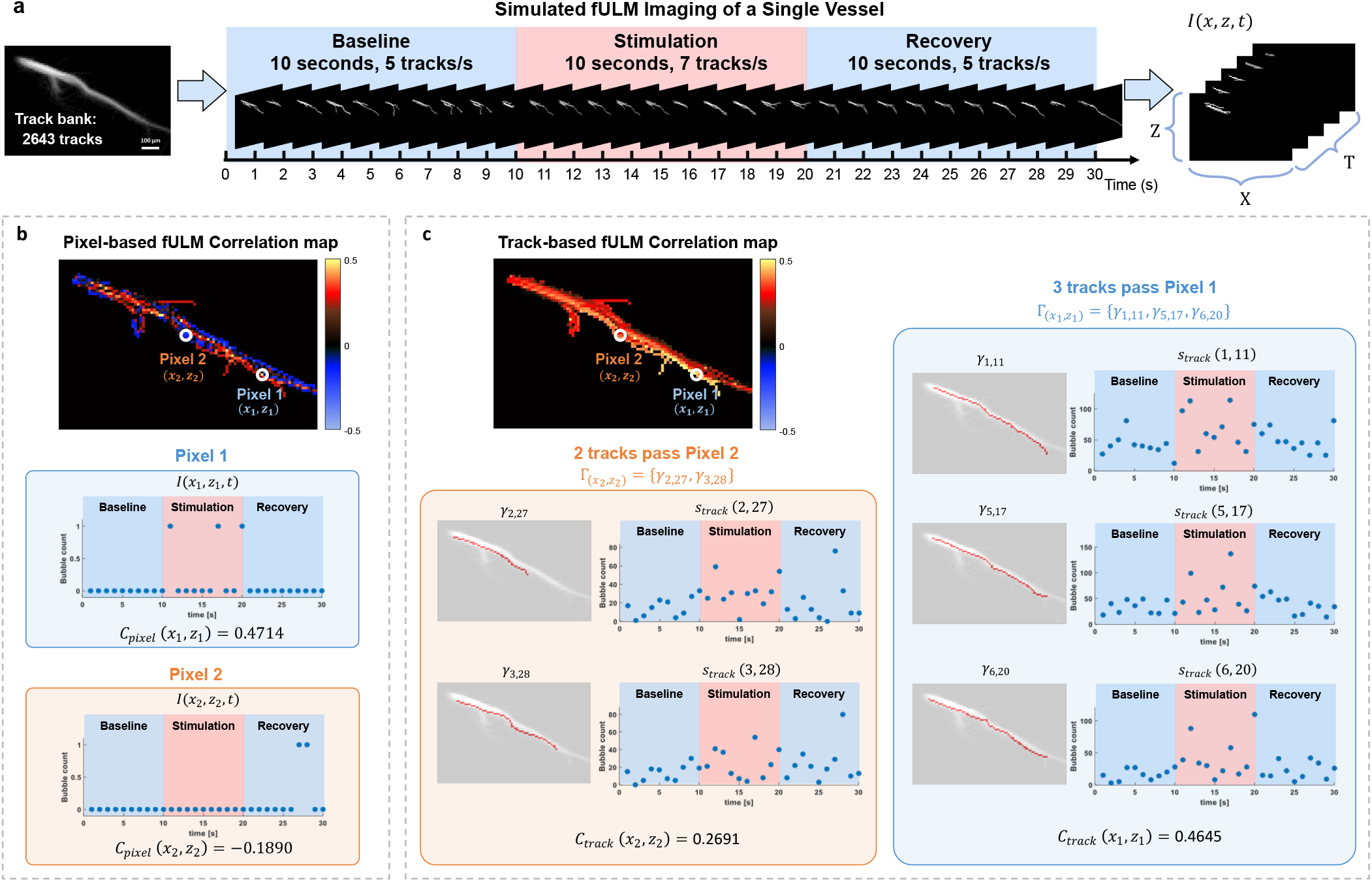
Illustration of pixel-based and track-based fULM methods using a simplified single-vessel simulation. (a) Generation of the simulation dataset. (b) Pixel-based fULM correlation map and temporal MB variations from two example pixels. (c) Track-based fULM correlation results, along with example MB tracks passing through two selected pixels and their associated temporal MB signal variations.

Upon completing ULM reconstruction, 2,643 unique MB tracks were identified within the single targeted vessel (Fig. 1a) and subsequently used as the track “bank” for simulation. A total of 30 ULM frames (T = 30 s) were simulated, with each frame representing one second of acquisition. The simulated data includes three phases: baseline (*t* ∈ [1,10] *s*), stimulation (*t* ∈ [11,20] *s*), and recovery (*t* ∈ [21,30] *s*). During baseline and recovery, blood flow was presumed to be stable, and 5 MB tracks were randomly sampled from the track bank per second. During stimulation, 7 MB tracks/s were selected to represent a 40% increase in blood flow. Note that the 40% increase is an artificially chosen value, used here solely to demonstrate a clear signal change in a small dataset for explaining the concept. In actual functional ultrasound contexts, the real hemodynamic response may be lower than 40%.

The number of MBs at any given frame *t* and pixel coordinates (*x, z*) is denoted as *I*(*x, z, t*). A stimulation pattern *r*(*t*) is introduced as a binary array of length T, with values of 0 during non-stimulation and 1 during stimulation:

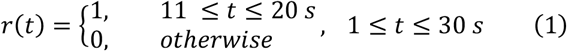

In conventional pixel-based fULM analysis, correlation calculation (e.g., Pearson’s) is typically conducted on the level of pixels:

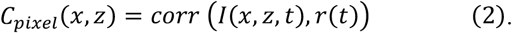

The resulting correlation map is shown in Fig. 1b.

In the proposed track-based method, each MB track is denoted by *γ*_*n,t*_, where *n* represents the track index. *γ*_*n,t*_ consists of a number of pixels: 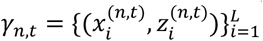, where *L* is the number of points in the track (track length).

Let Γ be a set of tracks and *N* represents the total number of tracks in Γ. For example, in Fig 1a, the set of all tracks over the 30 seconds in the simulation dataset can be defined as Γ_*total*_ = {*γ*_1,1_, …, *γ*_5,1_, *γ*_1,2_, …, *γ*_5,2_, …, *γ*_*n,t*_, …, *γ*_5,30_}, where corresponding *N*_*total*_ is 170.

For each pixel (*x, z*), a set of tracks that pass through the pixel is then defined as:

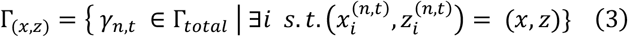

The track-based method shifts the focus of analyzing the correlation between MB count and the stimulus pattern from individual pixels to entire MB tracks, with the assumption that an individual MB track is the smallest responding unit to an invoked neural activity. This approach assumes that a single MB track within a single vessel segment should respond uniformly to the stimulation. The track-based approach is based on the fact that each MB track consists of data points acquired at millisecond-level intervals, and therefore all the MB events (i.e., MB count) along a given track is representative of the functional hemodynamic response within the vessel. As such, the signal values of all pixels along the track are aggregated, as a measurement of the collective activity in the vascular region where a MB traverses:

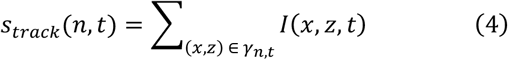

Once the *s*_*track*_(*n, t*) is computed, it is correlated with the stimulation pattern, producing a correlation coefficient for each track, denoted as *corr*(*s*_*track*_(*n, t*), *r*(*t*)).

This coefficient is then assigned to all pixels on the track. For any given pixel, the coefficients of all tracks passing through the pixel are averaged, resulting in a correlation value that more accurately reflects the response, as shown in Fig. 1c:

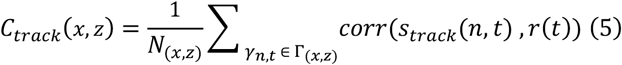

where *N*_(*x,z*)_ represent the total number of tracks that pass through pixel (*x, z*).

Assigning the same correlation value to all pixels along a track can be regarded as introducing temporal MB dynamic as a physiologically-informed prior, which leverages the assumption that adjacent pixel locations along the same MB track share similar hemodynamic responses. While this method may appear to reduce pixel-level sensitivity, this effect is mitigated by the fact that each pixel is traversed by a unique combination of tracks. As these contributions are averaged, the resulting map still retains the native spatial resolution of ULM. To further illustrate the principles with an example, in the pixel-based fULM correlation map (Fig. 1b), Pixel 1 has a relatively high correlation value (0.4714). During the non-stimulation period (0–10 seconds), no MB passed through this pixel. In the stimulation period (11–20 seconds), three MBs were detected, leading to signal fluctuations that align with the stimulation pattern. However, in Pixel 2, which is also an active pixel, no MB is detected during 0-26 s. Only two MB events occurred in the recovery period (27–28 s) which resulted in a negative correlation value. These random MB events obscure the estimation of blood flow variations, which is the key limitation of pixel-based fULM. In contrast, MB track-based correlation analysis reduces the influence of random fluctuations. For Pixel 1 (Fig. 1c), the track-based approach identified three tracks passing through the pixel 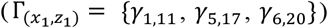. Summing the pixel signals along each track leads to elevated MB count during stimulation and positive correlation values.

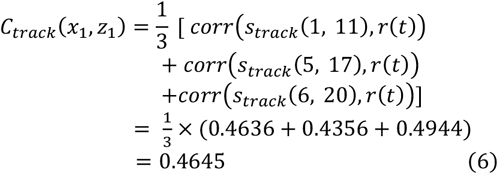

For Pixel 2, although the tracks passing through this pixel 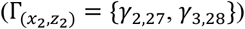 were detected during recovery period rather than the stimulation period, the aggregated signal along their entire tracks still exhibits a positive correlation with the stimulation pattern.

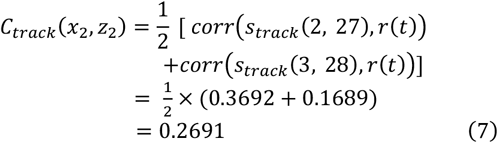

### Algorithm 1 MB Track-Based Algorithm

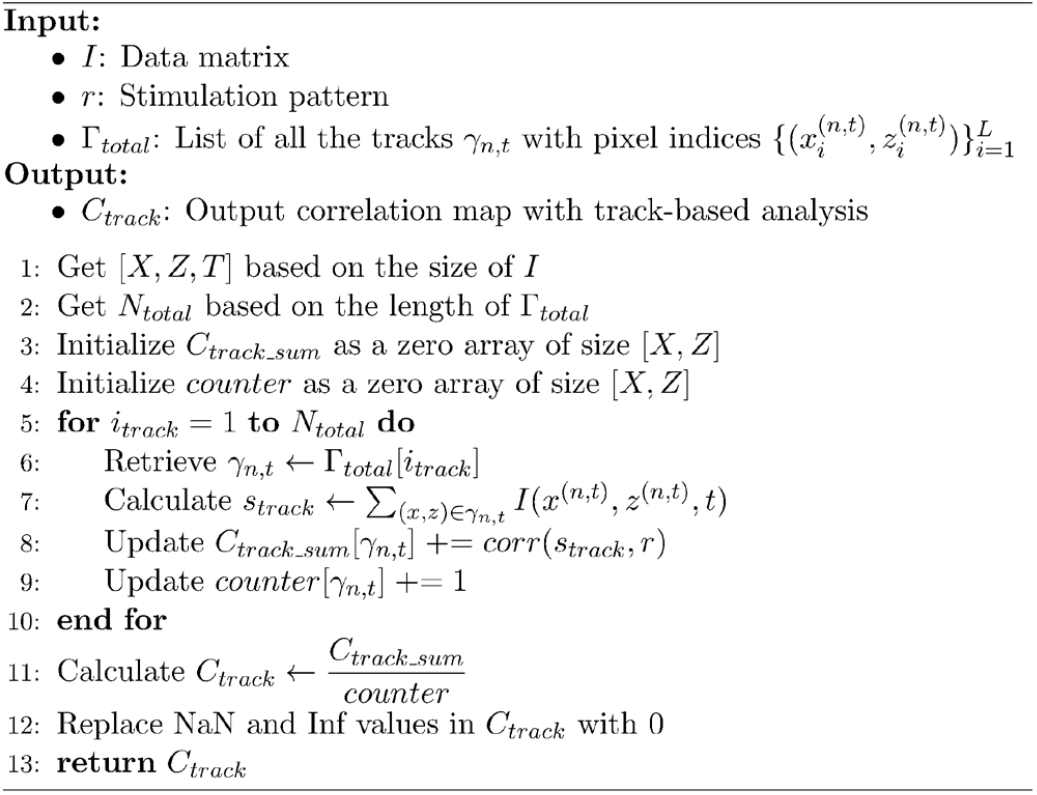

Unlike the scattered and fragmented patterns produced by pixel-based methods (Fig. 1b), the track-based map reflects the hemodynamics across the entire vessel segment (Fig. 1c), offering a more physiologically meaningful representation.

In practical implementation, it is unnecessary to explicitly search for all tracks passing through each pixel. To reduce computational complexity, we introduce an efficient algorithm by iterating over each track and incrementing the track count for each pixel it traverses. The pseudocode (Algorithm 1) illustrates the implementation. It initializes two zero vectors: *C*_*track*_*sum*_ to accumulate correlation values for each pixel and *counter* to count the number of tracks passing through each pixel, which is equivalent to *N*_*x,z*_ above. Finally, the averaged value for each pixel in the track-based correlation map is obtained by dividing *C*_*track*_*sum*_ by *counter*. This approach processes tracks sequentially, avoiding the need to explicitly indexing tracks with Γ_(*x,z*)_.

## III. Material and Methods

### A. In Silico Validation and Benchmarking

To quantitatively evaluate the performance of the track-based method, we designed a benchmarking framework that combines experimental and simulation data for analysis. In fULM experiments, it is inherently difficult to obtain physiologically verified ground truth activation maps, as the precise locations of activated microvasculature cannot be validated by any other *in vivo* imaging modalities. To address this challenge, we generated a set of *in silico* datasets, allowing us to benchmark different fULM analysis methods against the reference.

Fig. 2 explains the benchmarking process. The framework begins with a high-quality *in vivo* fULM dataset acquired from an anesthetized rat under whisker stimulation, which is referred to as the reference dataset. It consists of 15 stimulation cycles, each with a 15-second baseline, a 30-second stimulation, and a 15-second recovery period. From this dataset, a pixel-based correlation map was computed (Fig. 2a), where activation of the contralateral primary somatosensory cortex (S1) and ventral posteromedial nucleus (VPM) regions can be detected.

**Fig. 2.**
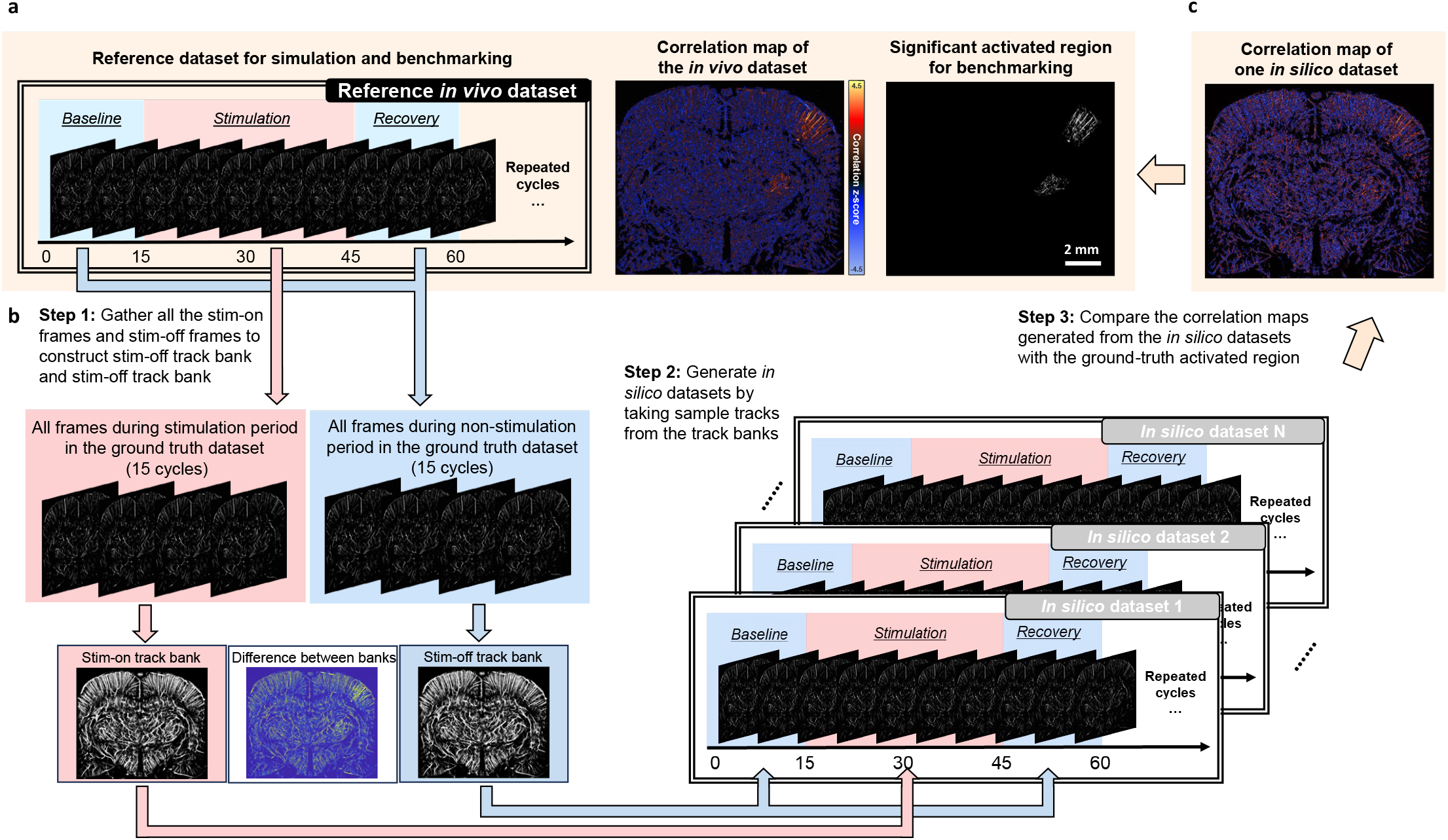
Pipeline for generating *in silico* datasets from a reference in vivo dataset and benchmarking them against a derived activation region. (a) A representative fULM dataset from an anesthetized rat under whisker stimulation is used as the reference dataset. A correlation map is computed over 15 stimulation cycles, and a significant activation region is defined by combining the significantly activated pixels with anatomical references to the primary somatosensory cortex (S1) and ventral posteromedial nucleus (VPM). (b) All frames during stimulation and non-stimulation periods are extracted from the in vivo dataset and grouped together to construct “stim-on” and “stim-off” track banks. These banks serve as sampling pools for generating in silico dataset. In silico datasets are constructed by randomly sampling tracks from the stim-on and stim-off banks to mimic controlled activation patterns. (c) The resulting correlation maps from these simulated datasets are then compared to the activation region defined in (a) to evaluate the performance of different analysis methods.

A benchmarking activation region was then defined by combining statistically significant pixels from the correlation map with anatomical priors from the Allen Brain Atlas [24] (S1 and VPM regions). Although this does not represent absolute physiological ground truth, it serves as a practical and consistent reference for benchmarking. To identify significantly activated pixels based on the corresponding p-value, each correlation value needs to be converted into a z-score using the Fisher’s transform, which has been used in previous studies [16], [25]. Here, a p-value of less than 0.05 was defined as significantly correlated, which corresponds to a threshold z-score of 2.8. By substituting n = 900 and *z*_*score*_ = 2.8 into the following equation:

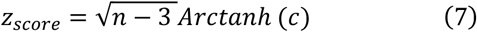

where n is the number of data points (900 acquisitions in total over the 15 cycles in this case) and *c* is the correlation coefficient, a correlation threshold for statistical significance can be determined. Pixels within the S1 and VPM regions exceeding this threshold were considered significantly activated (Fig. 2a).

*In silico* datasets were subsequently generated from the reference dataset, while preserving realistic vascular morphology and flow characteristics. The goal of the benchmarking process is to assess whether different fULM analysis methods can accurately recover the benchmark activation region from simulated datasets.

To generate the *in silico* datasets, two track banks were first constructed from the reference dataset (Step 1, Fig. 2b). All frames during stimulation periods across the 15 cycles were extracted and used to form the stim-on track bank, while frames from non-stimulation periods (baseline and recovery) were used to build the stim-off track bank. A subtraction between the two banks revealed a localized increase in track density in the right somatosensory cortex, indicating that stimulation-related changes in cerebral blood flow were effectively captured. These two track banks were subsequently used as physiologically-informed track pools for simulation.

In the second step (Step 2, Fig. 2b), *in silico* datasets were generated by taking sample tracks from the two banks at rates derived from the original *in vivo* dataset. Unlike the example in Section II, where MB track sampling rates were manually defined, here the average MB track production rates were directly estimated from the reference dataset: 2725 tracks/s during stimulation and 2705 tracks/s during non-stimulation, across 450 seconds of imaging. Tracks were randomly selected from the stim-on bank for stimulation periods and from the stim-off bank for non-stimulation periods, with the original temporal cycle structure preserved. To reduce variability due to random sampling, ten independent *in silico* datasets were created. For the next step, correlation maps from these simulated datasets (Fig. 2c) were compared against the benchmark activation region to evaluate analysis performance.

### B. Quantification of Sensitivity

To quantitatively validate that the track-based method achieves higher sensitivity and reduces the acquisition time, the receiver operating characteristic (ROC) curve was calculated across different number of cycles (e.g., 1, 3, 5, 8, 10, and 15 cycles) for each *in-silico* dataset. For each correlation map, a threshold was arbitrarily defined, with pixels exceeding this threshold considered activated. By changing the threshold, the sensitivity values at different specificity levels were assessed, generating an ROC curve. For any certain threshold value, a binarized activation map can be obtained and then compared with the benchmark region to calculate the true positive rate (TPR), true negative rate (TNR), false positive rate (FPR), and false negative rate (FNR). For example, the true positive rate (TPR), was calculated as the number of correctly classified activated pixels in the *in-silico* dataset at the selected threshold, divided by the total number of actual activated pixels. After generating the ROC curve, the area under the curve (AUC) was used to quantify overall sensitivity: the closer the AUC value is to 1, the more sensitive the method.

### C. Overview of animal imaging procedures

Female C57BL/6J mice aged 10-12 weeks were used for this study. All animal procedures were approved by the Institutional Animal Care and Use Committee at the University of Illinois Urbana-Champaign (IACUC Protocol number #22165). The overall experimental workflow consisted of following stages. Prior to surgery, all animals underwent a five-day handling to reduce stress during imaging. Each day, mice were gently hand-cupped for less than one minute and allowed to freely explore a custom 3D-printed body tube, which familiarized themselves with both the experimenter and the imaging setup [25]. After pre-surgical handling, mice were subject to survival surgery under anesthesia, during which both indwelling jugular vein catheterization and chronic cranial window surgery were performed (Fig. 3a).

**Fig. 3.**
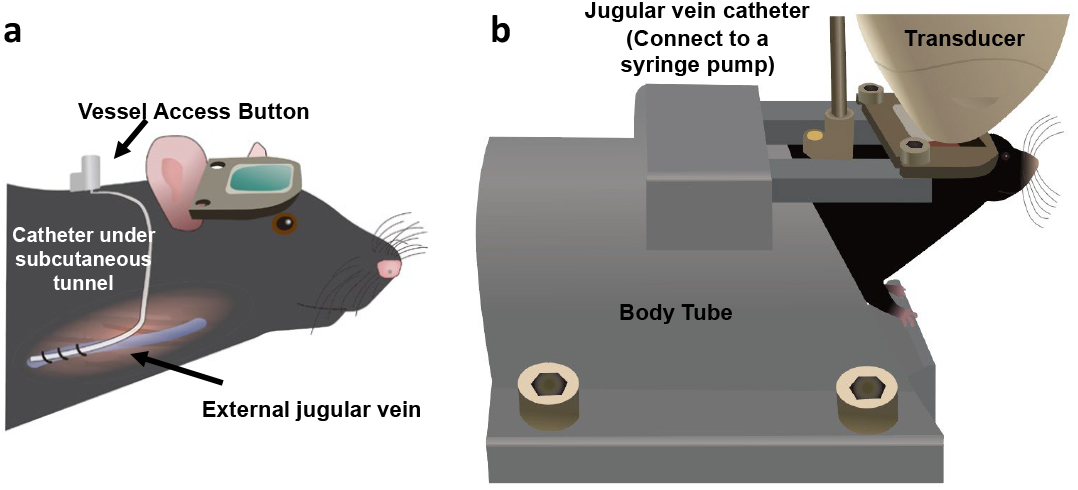
Schematic of the awake mouse fULM imaging setup. (a) Illustration of the indwelling jugular vein catheterization. (b) During imaging, the mouse is secured in a body tube, with the transducer placed on the headplate and a syringe pump connected to the Vessel Access Button.

The surgical protocol is detailed in the following sections. Postoperatively, animals were allowed to recover for one week. After recovery, mice were gradually habituated to be head-fixed through daily sessions in a body tube: 10 minutes on day one, 30 minutes on day two, and 60 minutes on day three [25], [26]. To compare fULM imaging between awake and anesthetized states, imaging was first conducted under awake conditions. Mice voluntarily entered the 3D-printed body tube and were secured in the head-fixation setup. The silicone rubber was gently removed, and ultrasound coupling gel was applied. The transducer was then positioned over the imaging plane at Bregma -1.8 mm. A syringe pump infusing Definity MBs at 40 μL/min was connected to the Vascular Access Button™ (VAB), and the fULM imaging session was initiated (Fig. 3b). Detailed imaging protocol are provided in later sections. After the awake imaging session, mice were anesthetized via intraperitoneal injection of ketamine (40 mg/kg body weight). Respiratory and heart rates were continuously monitored, and once physiological parameters stabilized—typically after 15 minutes—fULM imaging under anesthesia was performed following the same acquisition protocol.

### D. Animal Surgeries

On the day of surgery, the animal was anesthetized with isoflurane (3% for induction and 1%-1.5% for maintenance). A small incision was made rostral to the clavicle over the right jugular vein, which was then carefully isolated. Two pieces of sterile silk suture (6-0) (Oasis, MV-711-V) were used to ligate the vein, with one caudal piece near the clavicle and the other rostrally. A small incision was made in the jugular vein approximately 1 mm away from the rostral ligature. A catheter (Instech Laboratories, Inc., C20PU-MJV2011) prefilled with saline was gently inserted into the vein, and the caudal ligature was loosened to allow the catheter to advance toward the right atrium. After securing the catheter with the silk sutures, it was tunneled subcutaneously from the neck incision to an interscapular site, where a subcutaneous pocket was created with blunt dissection. The catheter was then trimmed to the appropriate length and connected to the VAB (Instech Laboratories, Inc., VABM1B/25). The incisions were sutured to secure the VAB in the subcutaneous pocket. The catheter was flushed with sterile heparin sodium (20 IU/ml) (Sagent Pharmaceuticals, NDC 25021-0400-30) immediately after the surgery and every following 3-5 days to maintain patency. After jugular vein catheterization, the mice were fixed in a stereotaxic frame. Cranial windows were prepared following established protocols [25], [26]. Dexamethasone (0.5 mg/kg) (Vet One, NDC 13985-533-03) was administered intraperitoneally to minimize edema. The scalp was incised, and the temporalis muscle was separated from the skull. A titanium headpost was secured to the skull using dental cement (Sun Medical Co., Ltd, Super-Bond C&B). A 4 mm × 8 mm region of the skull (Bregma 0 mm to Bregma -4 mm) was removed using a high-speed rotary micromotor (Foredom, K.1070). Subsequently, a transparent polymethyl pentene film (Goodfellow, ME311051) was placed over the brain and sealed with tissue adhesive (3M, 70200742529) and dental cement. The cranial window was protected with biocompatible silicone rubber (Smooth-on, Body Double-Fast Set).

### E. Image acquisition and processing

A Verasonics NXT ultrasound system (Verasonics Inc., Vantage NXT 256) was used in this experiment. A linear-array transducer (Verasonics Inc., L35-16vX) transmitting at a center frequency of 20 MHz was connected to the system. The transducer was mounted to a translation motor (Physik Instrumente, VT-80) using a 3D-printed holder, which allows for precise positioning of the imaged coronal plane. A plane wave compounding method using five steering angles (−2°, −1°, 0°, 1°, 2°) was implemented, yielding a post-compounded frame rate of 1000 Hz and an ensemble size of 800 frames per dataset. One dataset was acquired and stored every second. For an entire functional imaging session, data were collected across 8 stimulation cycles, each comprising 30 seconds of stimulation followed by 30 seconds of rest. During the stimulation periods, the system triggered an Arduino-controlled rod to tap the mouse’s whiskers.

The acquired radiofrequency (RF) signals were saved for subsequent offline reconstruction using the Verasonics build-in program, which produced the beamformed in-phase and quadrature (IQ) data with a pixel size of half the wavelength in axial direction and one wavelength in lateral direction. For each IQ dataset, a directional filter was applied to separate MBs moving toward and away from the transducer based on the positive and negative Doppler shifts [27]. Subsequently, singular value decomposition (SVD) filtering was performed to extracted MB signals from the tissue, with the cutoff adaptively determined to achieve objective and consistent filtering results [28]. The filtered data was then spline-interpolated to a pixel size of one-tenth of the wavelength (4.928 μm). Each interpolated frame was correlated with an empirically estimated point spread function (PSF) using 2D normalized cross-correlation. Local maxima in the correlation map were identified as MB centers. These coordinates were then linked across frames using the uTrack algorithm to reconstruct trajectories [29], [30], with only those persisting for at least 10 consecutive frames considered for valid MB tracks. The resulting tracks were subjected to either pixel-based or track-based fULM analysis, as described in Section II.

## IV. RESULTS

### A. In-silico Validation

Fig. 4 compares the results of two methods under different numbers of acquisition cycles. Fig. 4a shows correlation maps from one of the *in-silico* datasets. Since the *in-silico* datasets were sampled from the reference dataset, the resulting correlation maps are inherently limited in quality compared to those derived from the *in vivo* dataset. Nevertheless, the correlation maps still reveal activated S1 and VPM regions.

**Fig. 4.**
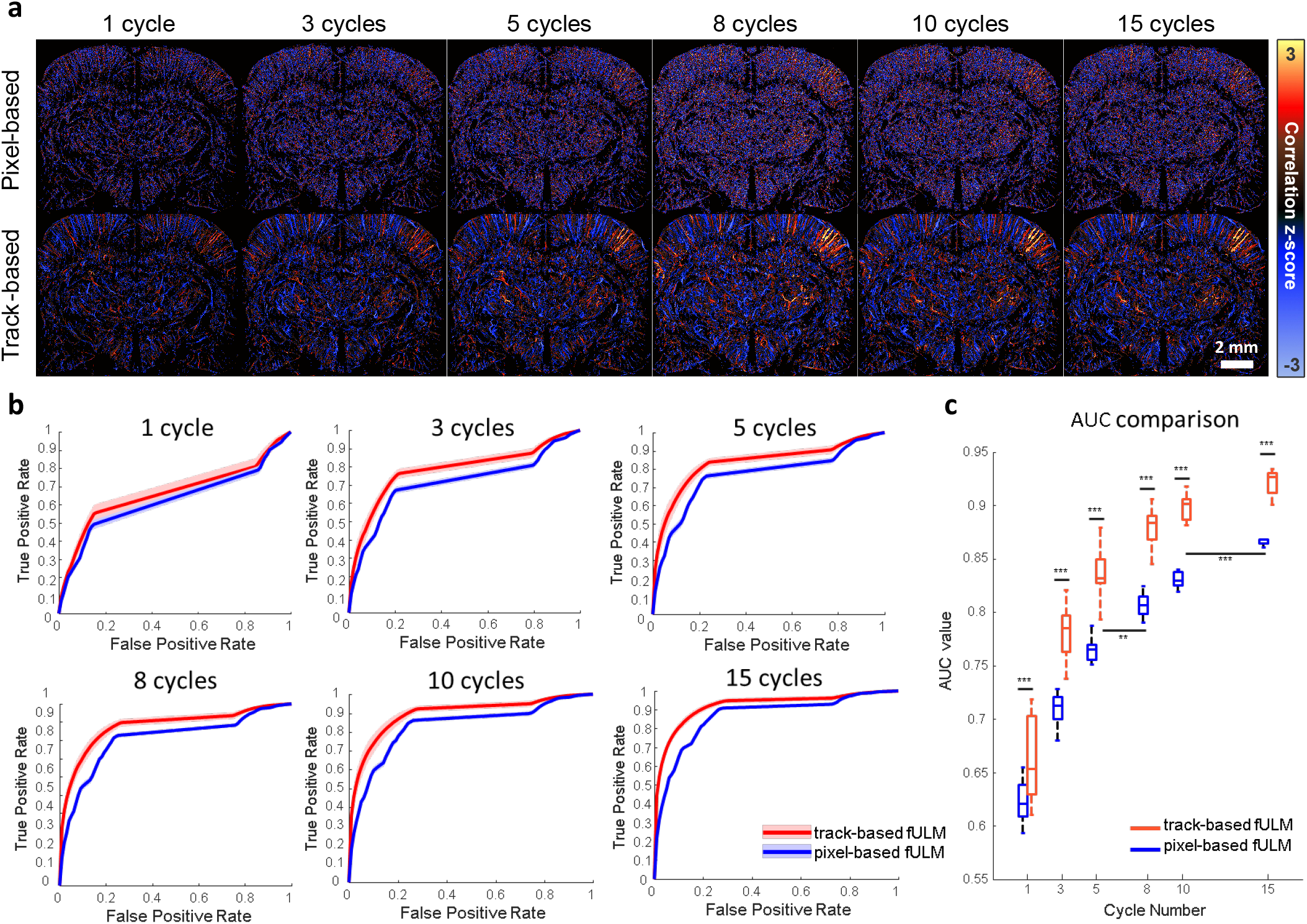
Validation results on the *in-silico* datasets. (a) Comparison between track-based and pixel-based analyses on one dataset under different stimulation cycles. (b) ROC curves of both methods across 10 datasets; solid lines indicate the average performance, and shaded areas represent inter-dataset variability. (c) Box plots of AUC values corresponding to the ROC curves in (b). (***, p <0.001; **, p<0.01)

Under the same correlation z-score, the track-based method reflects vascular activation more effectively and rapidly, with better contrast. Although some false positives in other brain regions make it difficult to distinguish the true S1 region at lower cycle number, as the number of cycles increases, the correlation strength in the activated regions becomes more pronounced, and false positives are progressively suppressed.

To reduce the influence of random variability from individual sampling instances, statistical analyses were performed on multiple *in-silico* datasets to assess overall performance (n = 10). In Fig. 4b, the ROC curve for the track-based method consistently lies above that of the pixel-based method, which higher sensitivity at the same specificity level. This conclusion is further supported by the t-tests of AUC values at the same cycle count in Fig. 4c.

The track-based method not only achieves better sensitivity under the same acquisition time but also reduces the number of stimulation cycles needed to match or exceed the performance of the pixel-based method, thus shortening data acquisition time. To validate this observation, ANOVA analysis was conducted on the AUC values for varying cycle numbers. For example, as shown in Fig. 4c, the track-based method achieves a significantly higher AUC value with five cycles than the pixel-based method with eight cycles. This demonstrates that the track-based method, as a post-processing technique, can identify activated regions more effectively with less experimental data.

### B. In-vivo Imaging

Fig. 5 compares the correlation maps obtained under anesthetized and awake conditions across different numbers of stimulation cycles. Results up to 8 cycles are shown here, because awake imaging inherently has higher functional sensitivity. It demonstrates that 8 cycles are already sufficient to reach the z-score corresponding to significant activation; therefore, it is unnecessary to acquire additional stimulation cycles as was done under anesthetized conditions. The overall findings are consistent with the *in-silico* dataset validation, where the pixel-based correlation method requires more cycles to detect the activated brain regions. In contrast, the track-based method can identify them with greater correlation z-score within the same number of cycles.

**Fig. 5.**
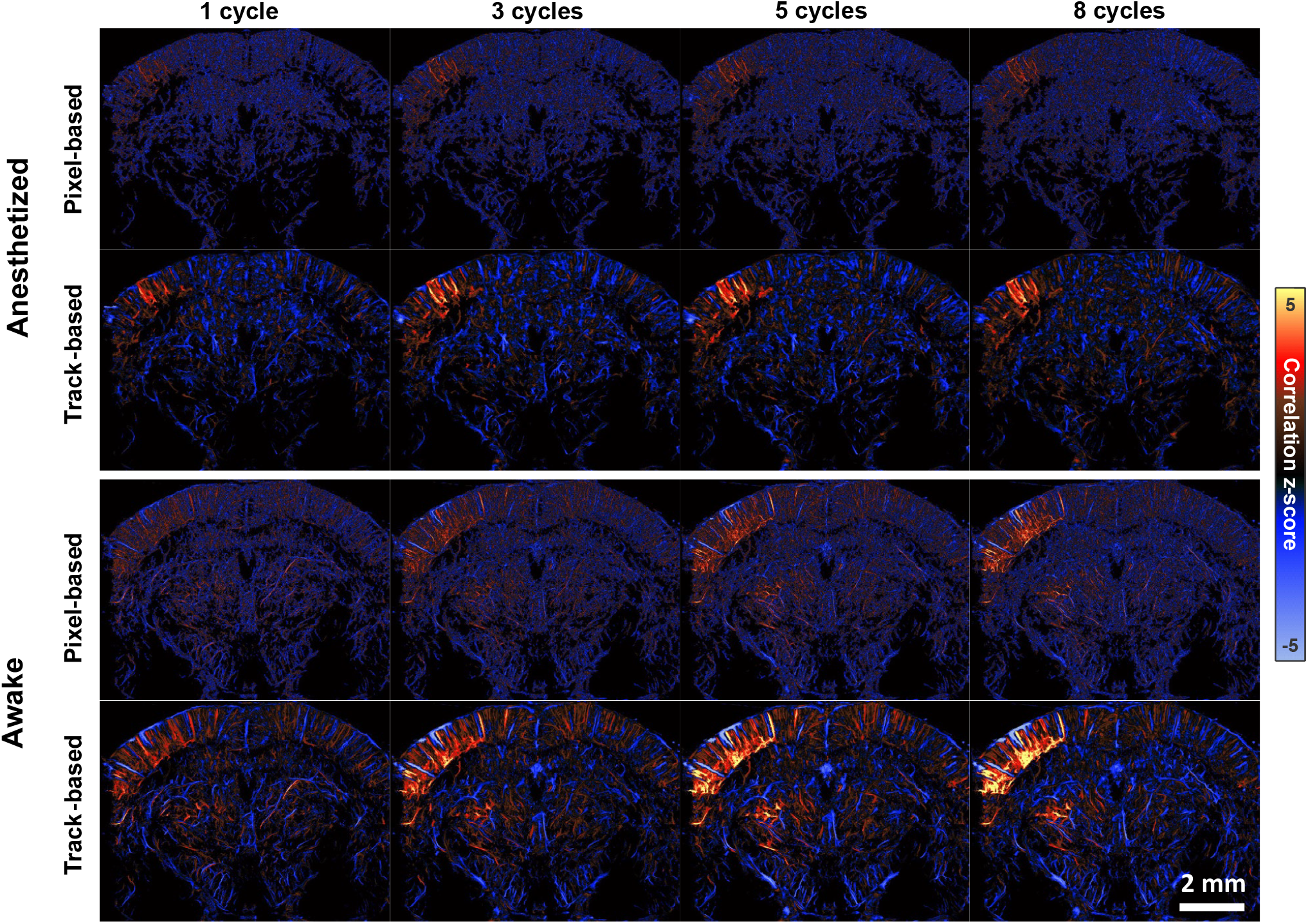
*In vivo* awake fULM results under different stimulation cycles. Track-based and pixel-based methods are compared across both anesthetized and awake states in the same mouse and imaging plane.

Under anesthesia, cortical responses were highly localized, with activation primarily restricted to the contralateral S1. Using the track-based fULM approach, this focal activation could be reliably detected within a single stimulation cycle, and the correlation strength increased progressively with the number of cycles. In contrast, much stronger and more spatially distributed responses can be observed in awake animals. Beyond the robust activation of S1, whisker stimulation in the awake state also indicated blood flow changes extending into the motor cortex, suggesting broader engagement of cortical networks. This may reflect the more complex functional connectivity present in the awake brain. Notably, thalamic responses—especially within the VPM—were more detectable in the awake state.

To further explore vascular specificity, we separately analyzed the upward and downward flow components, corresponding to venous and arterial flows in the cortex, respectively (Fig. 6). This classification scheme is particularly suitable for cortical penetrating vessels and has been employed in prior ULM/fULM studies as well [16], [26], [31]. While most arteries and veins in the activated S1 region exhibited strong positive correlations with the stimulus pattern, we identified one large vein (pointed by a green arrow in Fig. 6) that showed a negative correlation—an out-of-phase response in which blood flow decreased during stimulation and rebounded post-stimulation. This behavior, opposite to surrounding vessels, may result from passive regulation in downstream venous segments, where delayed hemodynamic responses are more likely to occur. Although further studies are needed to fully elucidate the mechanisms of such flow dynamics in the awake brain, our findings demonstrate the utility of the high spatial resolution of fULM in enabling vessel-specific hemodynamic observations. This capability offers unique insights into neurovascular response across subcortical regions, which are unattainable with conventional fUS.

**Fig. 6.**
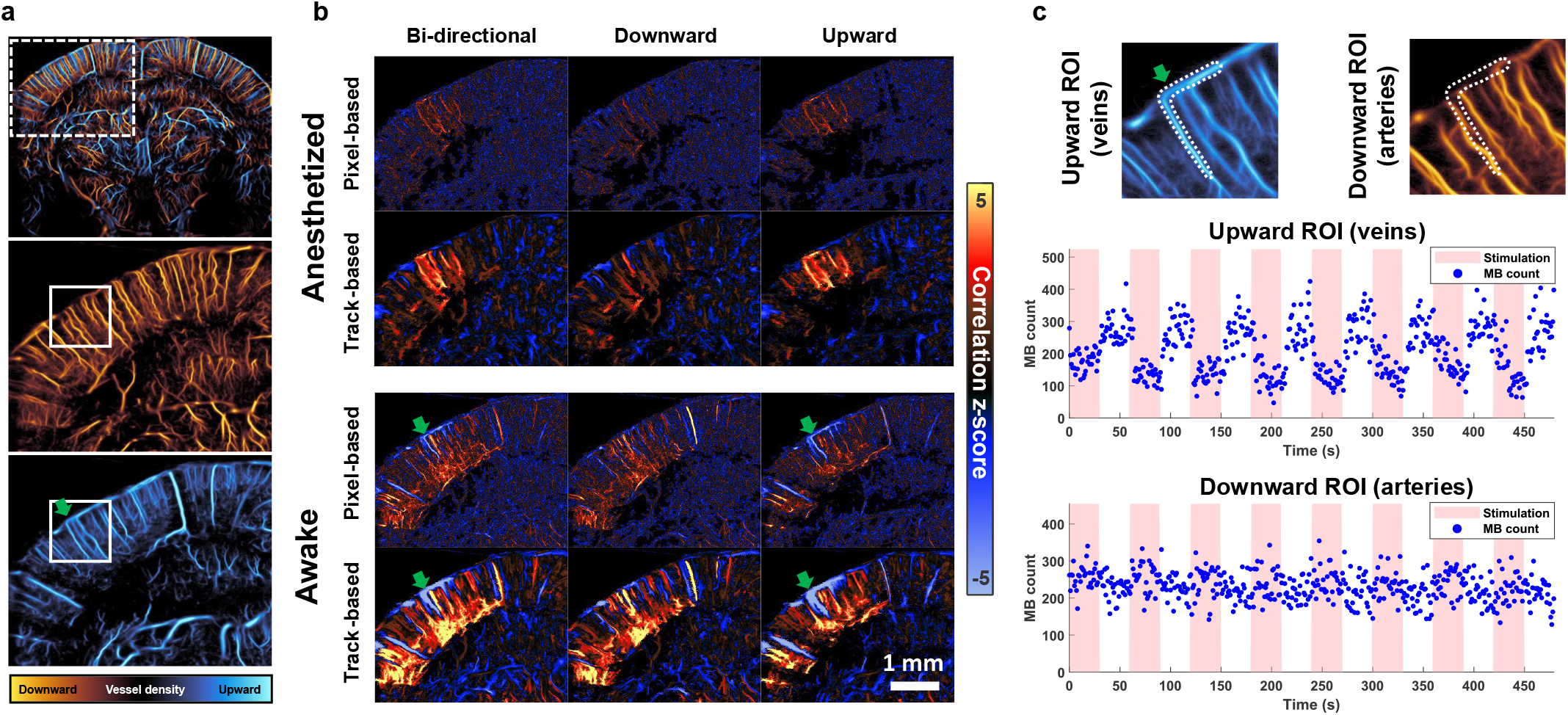
Arteriovenous response comparison within a cortical ROI. (a) A cortical subregion was extracted from the whole-brain image (white box) and separated into upward- and downward-flow datasets based on flow direction. (b) Pixel-based and track-based fULM results for bi-directional, downward, and upward flows under 8 stimulation cycles. (c) Temporal microbubble (MB) variations within a venous ROI (selected from the zoomed region in (a)) exhibit a negative correlation with the stimulation pattern. In contrast, the same ROI analyzed using the arterial-flow dataset shows a weak positive correlation.

## V. Discussion and Conclusion

This study introduces an MB track-based fULM method and its application in awake mouse functional brain imaging. To evaluate the effectiveness of the MB track-based method, a simulation study was performed, from which quantitative validation through ROC curve and AUC analyses validated the enhanced fULM performance. When applied to awake mouse imaging under whisker stimulation, the proposed method further demonstrated more robust performance over conventional fULM. Moreover, our findings revealed distinct responses of overlapping cortical arteries and veins, highlighting the benefits of the high spatial resolution enabled by fULM for capturing vessel-specific hemodynamic responses to neural stimulation that are inaccessible by conventional fUS (i.e., functional ultrasound imaging without using contrast MBs). Notably, even under anesthetized conditions, the proposed track-based method can still detect reliable brain activation mapping with only a few stimulation cycles.

Physiologically, the effectiveness of the MB track-based approach stems from the coherent nature of neurovascular responses within small vessel segment. Each MB track spans multiple, adjacent imaging pixels and captures the spatiotemporal continuity of blood flow within these imaging pixels. By integrating signals over MB tracks that are likely to exhibit similar hemodynamic responses within short time windows, it mitigates the sparsity problem (i.e., sparse MB count in time for each imaging pixel) of the conventional pixel-based method. Moreover, the dense population of MB tracks in ULM provides multiple independent observations for each pixel, which enables robust estimate of the response through averaging. Since the averaging is performed along the trajectories of the MB tracks, it does not introduce spatial blurring in the final fULM images. As a result, this track-based method enhances functional sensitivity without compromising spatial resolution of ULM. However, averaging along MB tracks can reduce the specificity in identifying the exact vessels responsible for hemodynamic changes. In practice, the length of MB tracks used for averaging can be adjusted to balance the trade-off between sensitivity in detecting neural responses and specificity in localizing the responding vessels.

Despite the advantages, the track-based method also has limitations. Its high sensitivity depends on adequate averaging over multiple MB tracks. However, when data is limited and individual pixels are not sufficiently averaged, random fluctuations in a single pixel can propagate along the entire track, increasing the risk of false positives and reducing specificity. As indicated by the AUC curves in Fig. 4c, track-based method shows much larger variation during the first cycle, but this variability rapidly diminishes as the number of cycles increases. This faster stabilization is due to the ability to collect and average tens of thousands of MB tracks within a single cycle, which enables rapid convergence toward the underlying vascular response. In contrast, pixel-based methods lack this internal averaging mechanism and thus require more acquisition cycles to reach similar stability. Nevertheless, sufficient pixel averaging is still essential for track-based method to get reliable activation mapping, especially in small vessels.

The proposed method could expand the potential applications of fULM in neuroscience. Traditional fULM applications have been limited by long acquisition time. By reducing the repetition of stimulation cycles, our MB track-based awake imaging could potentially enable broader and more practical applications in neuroscience community, particularly for applications focusing on deep brain regions where optical imaging techniques are difficult to reach. In addition, our awake fULM imaging setup also provides a more impactful and relevant tool for many neuroscience applications where anesthesia is not allowed (e.g., research in behavior). In the future, with the rise of wearable ultrasound imaging technologies [8], [9], further developments may achieve fULM in freely moving animals to support the investigation of brain activity under naturalistic conditions. Although intravenous MB infusion remains a challenge for imaging freely moving animals, the jugular vein catheterization method used in this study can enable stable MB infusion within a confined movement area, which lays the foundation for future fULM applications in freely moving animals.

Other future studies may include neurological validation to fully interpret fULM signals. Since fULM relies on neurovascular coupling, clarifying how *microvascular* dynamics reflect neural activity is critical for accurate interpretation. Characterizing the microvascular hemodynamic response function will help link fULM measurements to underlying neuronal processes, thereby enhancing the method’s relevance and reliability for neuroscience research.

## VI. Acknowledgement

We thank Dr. Qi You for the assistance with the experiment.

## Notes

This study was partially supported by the National Institute of Biomedical Imaging and Bioengineering and the National Institute of Neurological Disorders and Stroke of the National Institutes of Health under grant numbers R21EB030072 and R56NS131516, and by the National Science Foundation CAREER Award 2237166, and by the Chan Zuckerberg Initiative (CZI) Ben Berres Early Career Acceleration Award. The content is solely the responsibility of the authors and does not necessarily represent the official views of the NIH and NSF.

### Competing Interest Statement

The authors have declared no competing interest.

